# The Time-Course of Cancer Cachexia Onset Reveals Biphasic Transcriptional Disruptions in Female Skeletal Muscle Distinct from Males

**DOI:** 10.1101/2022.11.08.515691

**Authors:** Francielly Morena da Silva, Seongkyun Lim, Ana Regina Cabrera, Eleanor R. Schrems, Ronald G. Jones, Megan E. Rosa-Caldwell, Tyrone A. Washington, Kevin A. Murach, Nicholas P. Greene

## Abstract

**Background:** Cancer-cachexia (CC) is experienced by 80% of cancer patients, representing 40% of cancer-related deaths. Evidence suggests biological sex dimorphism is associated with CC. Assessments of the female transcriptome in CC are lacking and direct comparisons between biological sex are scarce. The purpose of this study was to define the time course of LLC-induced CC in females using transcriptomics, while directly comparing the effects of biological sex.

**Methods:** Eight-week-old female mice were injected with LLC cells (1×10^6^) or sterile PBS to the hind flank. Tumors developed for 1, 2, 3 or 4-weeks. Due to dimorphism between tumor weight in 3- and 4-weeks of development, these were reorganized as low-tumor weight (LT, tumor-weight ≤1.2g), or high-tumor weight (HT, tumor-weight ≥2g). Gastrocnemius muscle was collected for RNA-sequencing (RNA-seq). Differentially expressed genes (DEGs) were defined as FDR<0.05. Data were further compared to RNA-seq of male mice from a previous study.

**Results:** Global gene expression of female gastrocnemius muscle reveals consistent DEGs at all timepoints, all associated with type-II interferon signaling (FDR<0.05). Early transcriptomic upregulation of extracellular-matrix pathways was noted at 1wk (p<0.05), JAK-STAT pathway was upregulated in 2wk, LT, and HT. Type II interferon signaling was downregulated in 1wk, LT, and HT (p<0.05). A second major transcriptomic downregulation in oxidative phosphorylation, electron transport chain and TCA cycle were noted in cachectic (HT) muscle only (p<0.05). Male-female comparison of cachectic groups revealed 69% of DEGs were distinct between sex (FDR<0.05). Comparison of the top 10-up and down DEGs revealed downregulation of type-II Interferon genes was unique to female, while males show upregulation of interferon-signaling pathways.

**Conclusion:** We demonstrate biphasic disruptions in transcriptome of female LLC tumor-bearing mice: an early phase associated with ECM remodeling and a late phase, accompanied by onset of systemic cachexia, affecting overall skeletal muscle energy metabolism. Comparison of cachectic female-male mice reveals ~2/3 of DEGs are biological sex specific, providing evidence of dimorphic mechanisms of cachexia between sexes. Alterations to Type-II Interferon signaling appears specific to CC development in females, suggesting a new biological sex-specific marker of CC. Our data support biological sex dimorphisms in development of CC.

**Highlights:**

- While males show impairments in skeletal muscle energy metabolism in early stages of CC, early transcriptomic alterations impact ECM remodeling that precedes impairments in skeletal muscle energy metabolism in female tumor-bearing mice.
- 2/3 of differently expressed genes in skeletal muscle undergoing cachexia are biological sex specific.
- Downregulation of Type-II Interferon genes is unique to female mice, which displayed preserved gastrocnemius mass despite systemic cachexia, representing a potential therapeutic target for muscle mass maintenance in cancer-induced atrophy.
- Mechanisms of LLC-induced cachexia appear to be biological sex specific which needs to be considered in further study of mechanisms and therapeutic modalities.

## Introduction

Cancer cachexia (CC) is a syndrome experienced by up to 80% of cancer patients (1). Despite its first description in 1858, literature lacked a formal definition until 2011, when Fearon et al. (2) described cachexia as a multifactorial condition characterized by skeletal muscle mass loss, with or without fat mass loss, resistant to conventional nutritional support and leads to a progressive functional impairment (2, 3). CC can lower quality of life, reduce tolerance to anti-cancer drugs, and is directly responsible for 20-40% of cancer-related deaths (1, 4, 5). Despite the severity of this condition, mechanisms behind CC are not fully elucidated, and effective therapies are unavailable to cancer patients.

Even though biological sex plays an important role in health and various diseases (6–9) including CC, pre-clinical research is still male-dominant (10, 11). Transcriptomic analysis is a robust tool for uncovering etiology of disease (10, 12, 13). Yet, limited studies utilize this approach to identify transcriptomic alterations at the onset and initial development of CC (10, 12, 14–16), with fewer still considering female biological sex (10). Our group and others have recently shown biological sex differences during onset and progression of CC through time-course studies (10, 17–21), including muscle mass protection noted in females with systemic cachexia (17), but not in males. For instance, CC is known to be driven by systemic inflammation with characteristic induction of inflammatory cytokines including IL-6 and TNFα, followed by a causal sequence of a catabolic shift and loss of skeletal muscle mass. Male colorectal CC has long been described as IL-6 dependent, while more recent data suggest IL-6 independence in female colorectal CC (21–24). In addition, our laboratory reported early onset mitochondrial degeneration during development of LLC-induced CC in males (19), while such alterations are not present until development of cachexia in females (17). Combined current data suggest mechanisms of CC within a type of cancer are biological sex dependent, however the extent and identity of such differences remains largely unknown leaving a critical gap in the literature.

Here, we utilize RNA sequencing (RNA-seq) in a time-course fashion of tumor development to achieve the two-fold purpose of 1) understanding the transcriptomic profile of protected gastrocnemius muscle mass in female mice undergoing systemic LLC-induced CC, and 2) directly compare the gene expression landscape between males in females during the time course of early cancer development. About two-thirds of all dysregulated genes were distinct between sexes, demonstrating unique biological sex signatures and thus likely biological sex specific mechanisms in LLC-induced CC. Sex-specific considerations are therefore imperative for developing individualized therapeutics for improving muscle mass during cancer.

## Materials and methods

### Animal Interventions

Female C57BL6/J mice (Jackson Laboratories, Bar Harbor, ME; *n* = 40) from a larger study were randomly selected (17). Phenotypic statistics for this subset were performed to validate representation of the larger dataset. Animals were kept on a 12:12-h light-dark cycle and given access to normal rodent chow and water for the duration of the study. All animal protocols were approved by the University of Arkansas Institutional Animal Care and Use Committee, in accordance with the ethical standards (1964 Declaration of Helsinki).

#### Lewis Lung Carcinoma allograft

Lewis lung carcinoma (LLC) cells (ATCC, CRL-1642) were grown as described (17). At eight-weeks of age, female mice were subcutaneously injected with either LLC cells (1 × 10^6^) suspended in 100μL PBS or equal volume sterile PBS (control) to the right hind flank. Tumors were allowed to develop for 1, 2, 3, or 4-weeks; 4-weeks is a timepoint commonly associated with mild cachexia in this model, majorly utilizing male subjects (18, 19). As we noticed a dichotomous pattern in tumor weight, 3 and 4wk mice were regrouped into low (LT, tumor-weight ≤1.2g) and high (HT, tumor-weight ≥2g) tumor bearing (17) n=8/group. Control (PBS) mice were age-matched with 12-weeks-old mice.

### Tissue collection

Mice were anesthetized with isoflurane prior to euthanasia and tissue wet weight of gastrocnemius, plantaris, soleus, extensor digitorum longus (EDL), tibialis anterior muscles of both limbs, along with heart, spleen, liver, and gonadal fat were assessed. Samples were snap-frozen in liquid nitrogen and stored at −80°C for further utilization.

### RNA isolation and quality check

Total RNA was extracted as described (12). Gastrocnemius was selected due to its heterogeneous fiber type composition, and to address biological sex differences on CC onset and development by allowing comparisons to our previous study in males (12). Total RNA concentration and purity were determined using BioTek Take3 micro-volume microplate with a BioTek Synergy HTX multi-mode plate reader (BioTek Instruments Inc., Winooski, VT) and 260/280nm ratios and RNA concentrations were obtained. Samples were used if 260/280 ratios were of acceptable (>2.0) quality.

### RNA Sequencing and data analysis

Complete data output and analysis can be found online for all RNA-sequencing and Pathway analysis associated data in the *Supporting Information*.

RNA-sequencing of gastrocnemius muscle was performed by the genomics core at Michigan State University. Briefly, libraries were prepared using Illumina TruSeq Stranded mRNA Library Preparation Kit (Illumina, San Diego, CA) per manufacturer recommendations. Libraries were divided into pools for multiplex sequencing using Illumina HiSeq 4000 flow cell in a 1×50bp single read format. Base calling was completed by Illumina Real Time Analysis (RTA) v2.7.7, output was demultiplexed and converted to FASTQ files with Illumina Bcl2fastq v2.19.1. FASTQ files were organized by assigning each sample to designated groups according to condition and timepoint (PBS, 1wk, 2wk, LT, HT). Pre-alignment QA/QC read was performed to assure quality, followed by a STAR 2.7.8a index alignment – assembly: *mus musculus* (mouse) – mm39. QA/QC was repeated post alignment. Normalization was performed by using Median ratio for DESeq2 in Partek. A filter of 30 minimum reads across all samples was applied. DESeq2 analysis with comparisons of each timepoint (1wk, 2wk, LT, HT) against control (PBS) and LT against HT was performed. Results were downloaded in text format, and further organized in Excel. Cutoffs for DEGs (differentially expressed genes) were performed at 0.05 false discovery rate (FDR-adj. P-Value). Pathway analysis was performed on DEGs for each comparison through G:profiler (25) with following settings: all results, with statistical scope considering only annotated genes, significance threshold of g:SC threshold (0.05) and numeric IDs treated as ENTREZGene_ACC.

Data used for analysis can be found in Supporting Information S1, Supporting Information S2, Supporting Information S3, Supporting Information S4, Supporting Information S5, Supporting Information S6, Supporting Information S7, Supporting Information S8, and described in Supporting Information Description.

### Comparison of RNA sequencing to MitoCarta

DEGs were cross-referenced with the Mouse MitoCarta 3.0 (26). Cross-reference was conducted by utilizing a custom computer software provided by Kevin B. Greene.

### Comparison of RNA sequencing to male data set

Comparison of RNA-seq data from the present study with a subset of male mice of a previous study from our group (12) was performed to assess biological sex differences in response to CC. FASTQ files from male mice were uploaded to Partek, and analysis was conducted with the same parameters utilized for female mice analysis as above. Cross-referencing of DEGs of female and male was performed by custom R command provided by Dr. Aaron Caldwell. Further DESeq2 analysis was conducted on Partek by directly comparing female and male mice RNA-Seq data with the same parameters described, specifically a 2X2 comparison of biological sex (male v female) to cachexia (PBS v cachectic (4wk [male]/HT [female]).

### Statistics

For phenotypic data, a one-way ANOVA was utilized for each dependent variable. When significant F-ratios were found, differences among means were determined by Tukey’s post hoc test. For all statistical tests, the comparison-wise error (α) rate of 0.05 was adopted. Data were analyzed through GraphPad Prism (La Jolla, CA, USA). Data expressed as mean ± standard error of the mean (SEM). Data from RNA Sequencing analysis was analyzed using DESeq2 through Partek Flow. False discovery rate (FDR, reported FDR step-up adjusted P-value) was controlled at < 0.05.

## Results

### Phenotypic description of LLC-induced muscle atrophy across time course

Phenotypic analyses were performed on a cohort of animals from a larger study (17) (n = 8/condition). Tumor-free body weight was not different between experimental groups compared to PBS. EDL mass was not different between experimental conditions. Plantaris showed ~11% higher mass in LT compared to HT, but no differences in cancer groups when compared to PBS control (Table1, p=0.04). Soleus and tibialis anterior muscles where 19.3% and 9.3% lower in HT compared to PBS, respectively (Table 1, p=0.002, and p=0.04). Gastrocnemius muscle of tumor bearing groups was not statistically different from PBS. As for visceral organs, liver mass of LT and HT were 23% and 43% higher than PBS (Table 1, p=0.02, p<0.0001). Spleen mass was higher in HT by 26.5%, 35%, 48.5%, 44%, compared to 1wk, 2wk, LT, and PBS, respectively (p<0.0001). Gonadal fat was 55.9% lower in HT compared to PBS (Table 1, p=0.04). Phenotypic data demonstrate multiple hallmarks of CC in HT, confirming induction of LLC-induced CC for the current selected subset.

**Table 1.** Phenotypic Data of Subset. Data expressed as mean +− SEM. P<0.05. Different letters indicate statistical significance (p<0.05).

| Group | PBS | 1wk | 2wk | LT | HT |
| --- | --- | --- | --- | --- | --- |
| Body weight (g) | 18.89± 0.21ab | 17.64± 0.18a | 18.61 ± 0.20a | 19.91± 0.44b | 21.24± 0.57 |
| Tumor Weight (mg) | N/Aa | 33.85 ±4.83a | 220.2±41.7ab | 506.4 ±106.4b | 3078±141 |
| Body Weight – Tumor Weight (g) | 18.89± 0.21ab | 17.60±0.19b | 18.39 ± 0.23ab | 19.41±0.38a | 18.16±0.54 |
| Gastrocnemius (mg) | 89.60±1.17abc | 85.42±1.88ac | 88.16 ±1.45abc | 94.63 ± 1.65b | 85.58±2.17 |
| Soleus (mg) | 7.3±0.18a | 6.43±0.18ab | 6.84±0.35ab | 7.28±0.17a | 5.89±0.27 |
| Plantaris (mg) | 12.78±0.43ab | 12.43±0.36ab | 12.65±0.28ab | 13.38±0.44a | 11.88±0.28 |
| EDL (mg) | 8.00±0.40 | 7.75±0.13 | 7.88±0.24 | 8.18±0.24 | 7.28±0.18 |
| TA (mg) | 36.03±1.20a | 35.27±1.23ab | 35.76±0.80ab | 36.31±0.86a | 31.99±0.83 |
| Fat (mg) | 313.60±32.41a | 238.80±43.78ab | 270.10±23.88ab | 340.00±35.25a | 175.40±30 |
| Heart (mg) | 102.9±3.32 | 100.80±1.80 | 102.40±4.8 | 105.30±2.98 | 105.20±3. |
| Liver (mg) | 729.30±31.08a | 804.30±31.68ab | 825.40±45.73ab | 896.50±45.73bc | 1043.00±47 |
| Spleen (mg) | 78.93±4.86a | 67.14±3.75a | 95.86±7.99a | 118.1±5.81a | 328.20±28 |

### Global gene expression analysis showed substantial transcriptome shift in high-tumor group only

To identify muscle transcriptome shifts in CC, we performed RNA-seq of gastrocnemius muscle at all timepoints. We identified 2,958 upregulated and 2,052 downregulated DEGs across all time points (Figure 1a, adj. P-Value<0.05). Most DEGs were in the HT group (83% of total DEGs) (Figure 1b-e adj. P-Value<0.05). DESeq2 analysis showed 2,446 upregulated, and 1,856 downregulated DEGs in HT compared to PBS while all the other timepoints combined showed 512 up- and 195 downregulated DEGs (Figure 1b-e, adj. P-Value<0.05). Comparing LT against HT we noted 1,000-up and 1,343-downregulated DEGs (Figure 1f, adj. P-Value<0.05). The top ten up- and downregulated genes of each timepoint are in Figure 2a-d. The top ten upregulated genes showed an elevation in genes associated with antioxidant action (*Mt3, Mt2*) along with cell structure associated genes (*Lgals3, Adam12*) in 1wk mice. Interestingly, various Guanylate Binding Proteins (*Gbp2*, *Gbp3, Gbp4 Gbp5*) were downregulated in 1wk mice and remained downregulated throughout later timepoints along with other immunity and interferon-associated genes (*Entpd1, Gcnt2, Atp8b1, Trim21 Wars, Irf1, Nlrc5, Cd274)* (Figure 2a-d).

**Figure 1.**
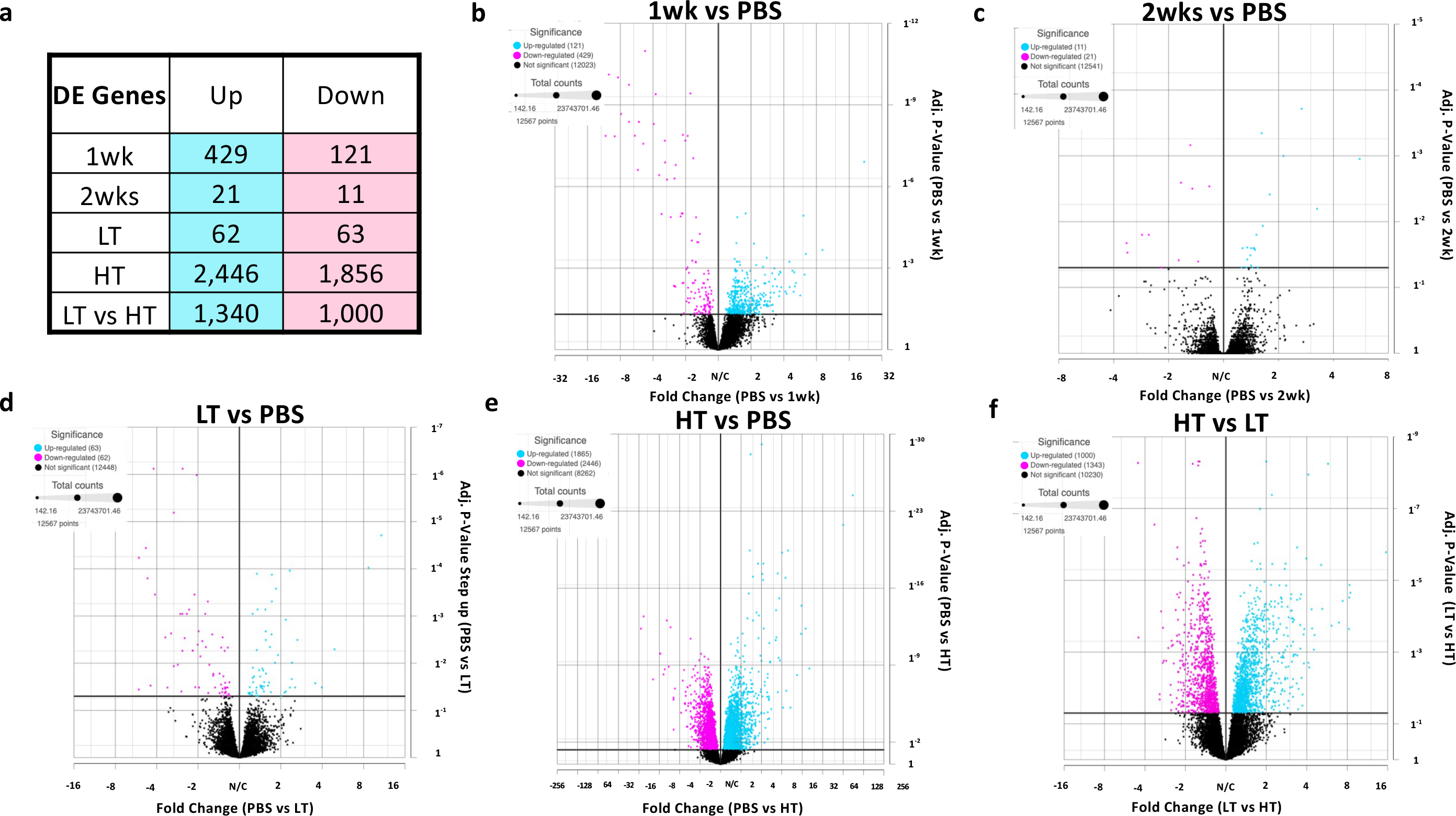
(a) Total number of Differentially Expressed (DE) Genes compared to PBS control. LT vs HT comparison was added. Volcano Plots for 1wk (b), 2wk (c), LT (d), HT (e), compared to PBS. HT compared to LT (F) was added. FDR<0.05.

**Figure 2.**
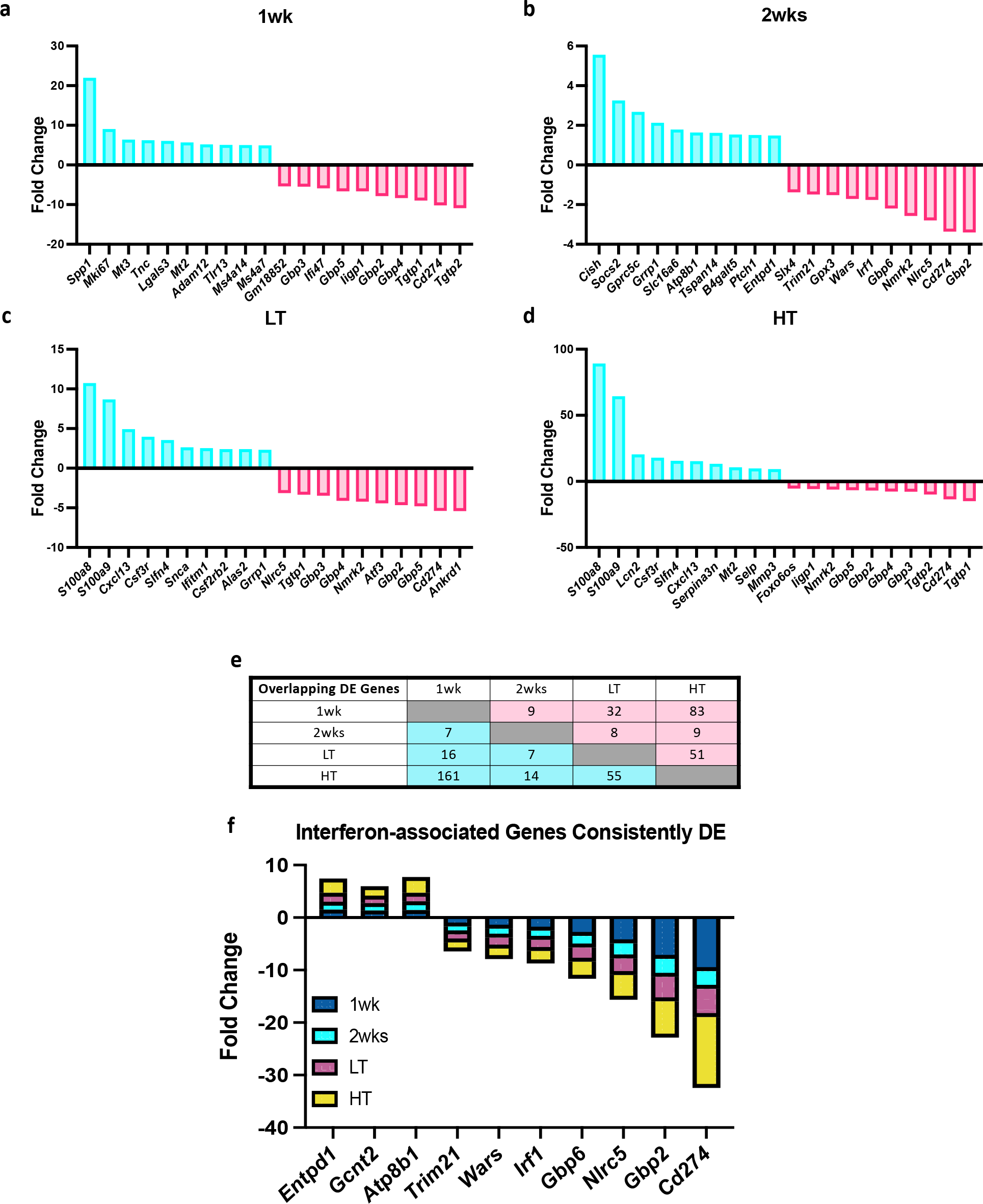
Top 10 Up- and Downregulated DE Genes. Tumor-bearing 1 week (1wk) (a), 2 week Low Tumor (LT) (c), and High Tumor (HT) (d) compared to PBS control. Total number of matches of up and down-regulated Differentially Expressed (DE) Genes compared to PBS control across all comparisons, LT vs HT comparison was added (e). Consistently DE genes across all cancer time points are associated with interferons (f). FDR<0.05.

To investigate differences and similarities in DEGs across timepoints, we investigated distinction in DEGs across timepoints (Figure 2 e-f). Overlapping DEGs between tumor-bearing comparisons found in Figure 2e. We subsequently identified 10 overlapping DEGs (3 upregulated and 7 downregulated) in all tumor-bearing groups compared to control (Figure 2f). Overlapping genes were associated with immune response, specifically Type-II-interferon signaling.

### Pathway analysis revealed biphasic biological alterations in CC development

Next, we explored functional ontology of the altered gene networks. The top up- and downregulated pathways (up to five) of KEGG, Reactome, and WikiPathways biomolecular pathways libraries are shown in Figure 3a-b. Phagosome associated pathways were amongst the upregulated pathways in 1wk group, along with multiple pathways suggesting ECM remodeling, changes in cell structure and oxidative stress response. At two weeks after tumor introduction, the JAK-STAT pathway was the only significantly upregulated pathway (not shown), which remained significant in the LT group, along with cytokine-cytokine receptor interaction pathway. In the HT group we identified upregulation of Proteasome, mRNA processing, Translation processing, and multiple inflammation associated pathways. Multiple immune system-related pathways were downregulated at 1wk along with type-II-interferon pathways, also consistently dysregulated in LT and HT groups. The most downregulated pathways in HT were associated with metabolism and mitochondrial systems, including oxidative phosphorylation (OXPHOS), ATP synthesis, TCA cycle, and Electron transport chain (ETC) pathways.

**Figure 3.**
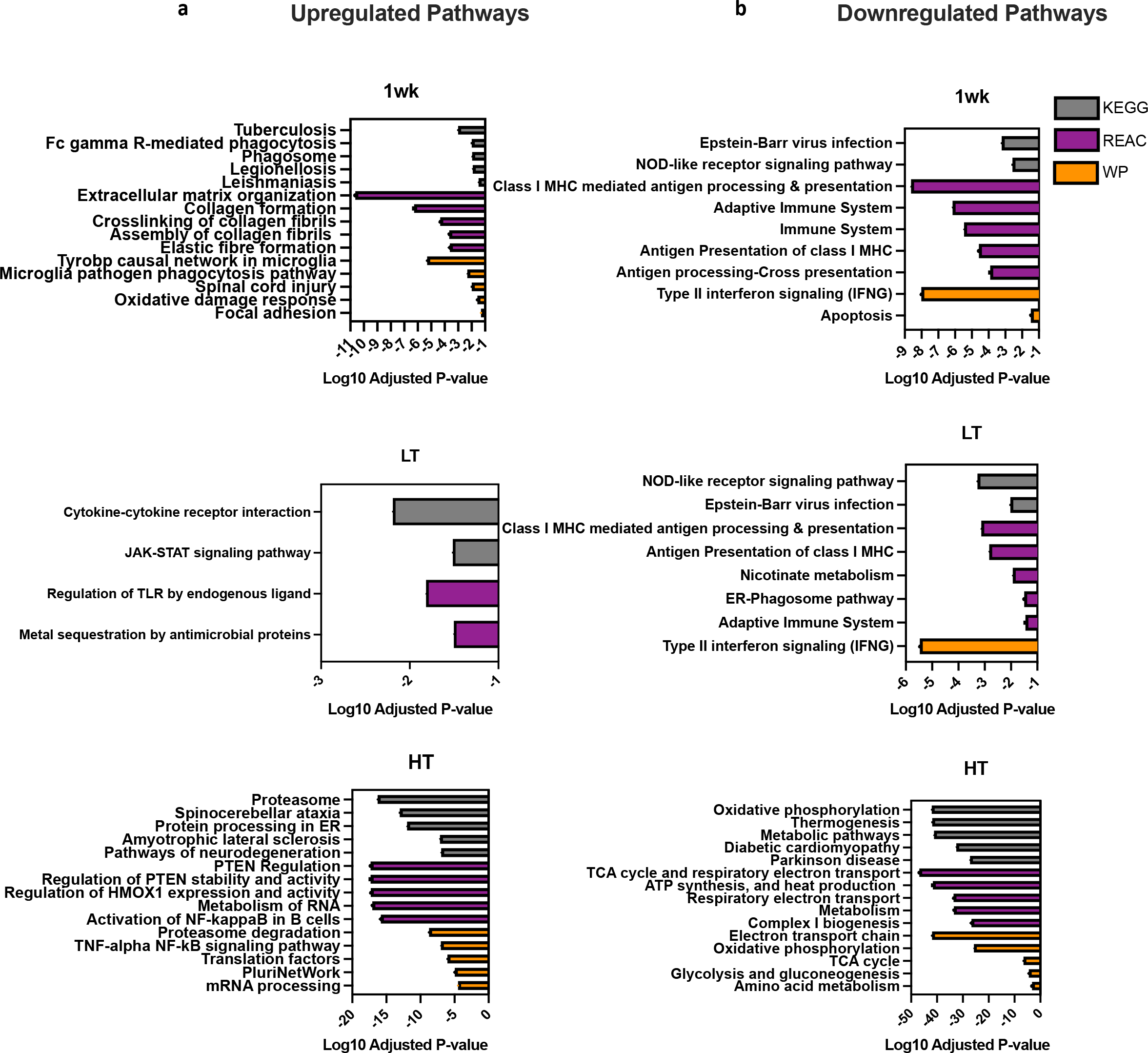
Top 5 Up-(a) and Downregulated Pathways (b) for KEGG Reactome, and WikiPathways. Tumor-bearing 1 week (1wk), Low Tumor (LT), and High Tumor (HT) compared to PBS control. Adjusted P-value<0.05.

### Comparison of RNA sequencing to MitoCarta shows striking alterations to mitochondria-associated genes in CC

Considering substantial disruption of mitochondria-associated pathways, we compared the current dataset with MitoCarta 3.0, composed of 1,141 known mitochondrial-associated genes (26). Altogether, overlapping DEGs represent 6% and 41% of MitoCarta genes that were up- and downregulated with LLC, respectively (Figure 4a-b). Very few genes matched the MitoCarta in the pre-cachectic groups. In the HT group, 55 up- and 421 downregulated genes were matched (Figure 4c). The top 20 MitoCarta matched are shown in Figure 5d including *Alas2, Bnip3*, and *Casp8* (Figure 4d). *Cox5a*, a subunit of Cytochrome C Oxidase and multiple mitochondrial transporters genes were amongst the top 20 downregulated genes, including *Slc25a47, Slc25a25*, and other *Slc* genes (Figure 4d).

**Figure 4.**
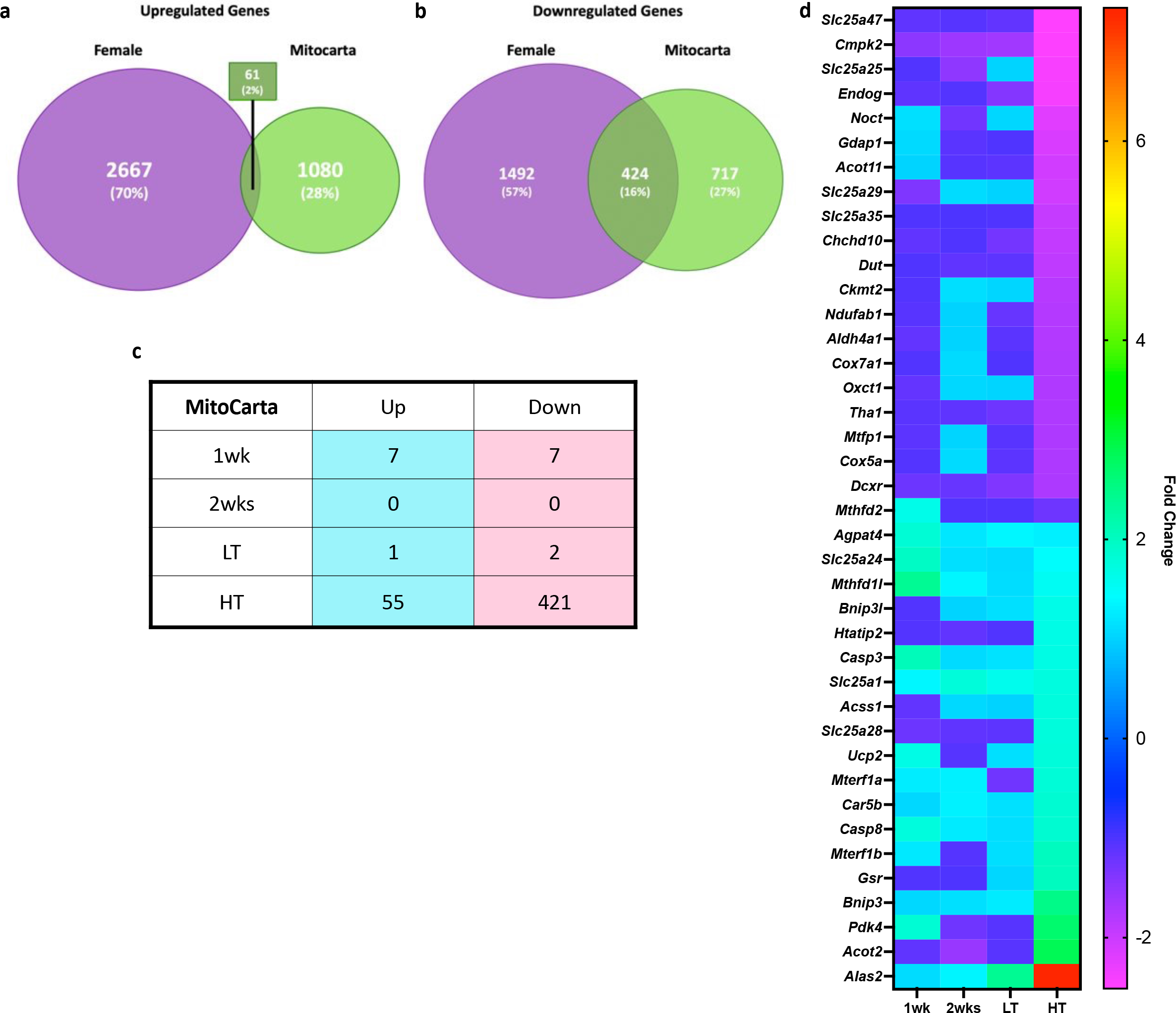
GE Genes Tumor-bearing Female and MitoCarta 3.0 Cross-reference. Venn Diagram of Upregulated Genes (a), Venn Diagram of Downregulated Genes (b), Matching Genes (c) of Female and MitoCarta 3.0 datasets, Top 20 Up- and Downregulated matching Genes of Tumor-bearing Female and MitoCarta 3.0 (d) genes organized from lower to higher FC in HT.

**Figure 5.**
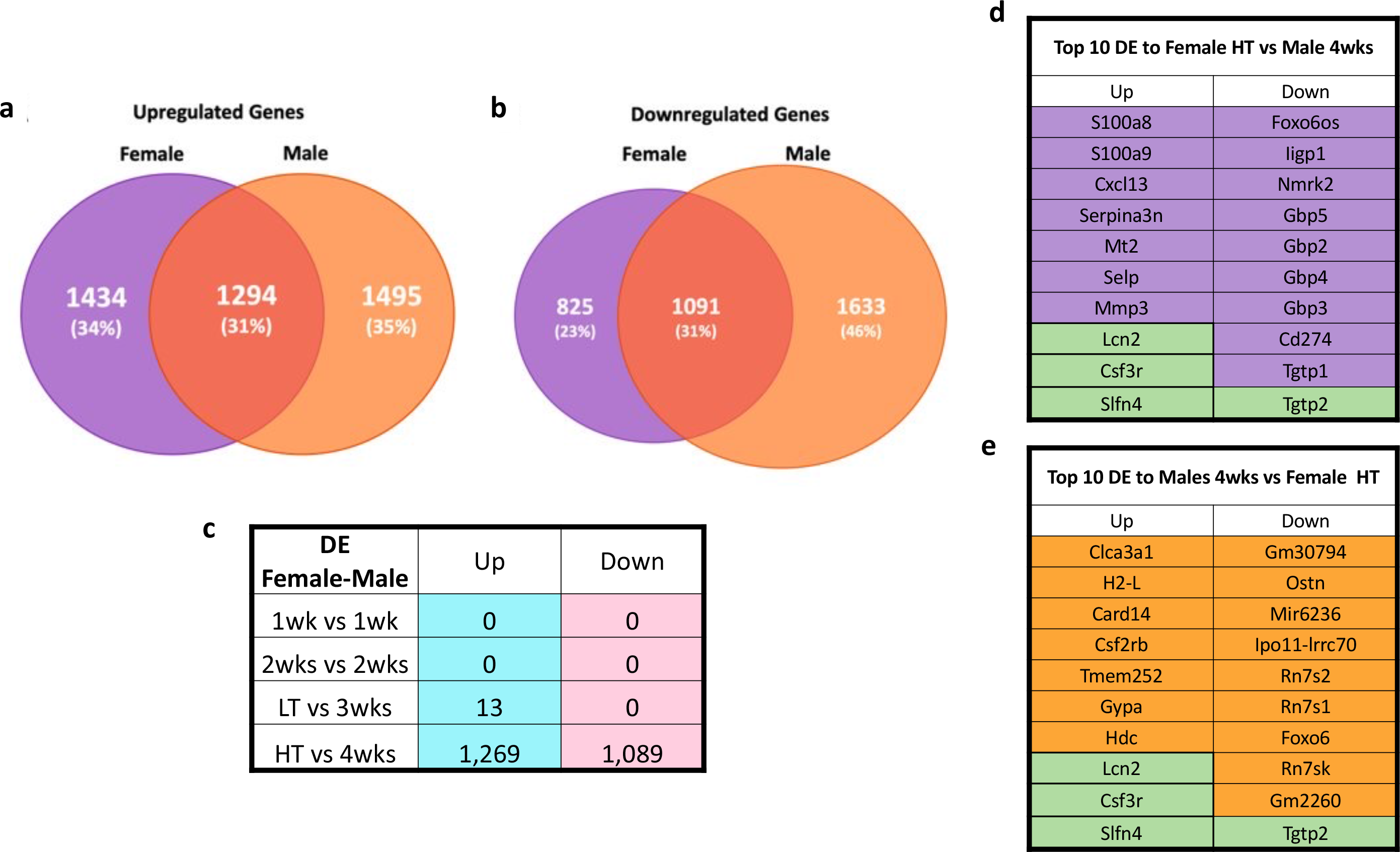
Female and Male Comparison. Venn Diagram of Upregulated Genes (a), Downregulated Genes (b), Matching Genes (c) of Female and Male datasets, Top 10 Genes Female HT vs 4wks (d), Top 10 Genes Male 4wks vs Female HT. Purple cells= genes unique to females, orange cells= genes unique to males, and green cells= common genes between biological sexes. FDR<0.05

### Comparison of RNA sequencing to males revealed biological sex dimorphism in transcriptomic alteration in CC

As biological sex plays an important role in CC (10, 17, 18), we cross-referenced female data with our previously published study utilizing LLC-induced cachexia in males (12). We previously reported phenotypic data for this male subset including an ~17% lower gastrocnemius mass in cachectic animals compared to PBS control (12). For consistency, we reanalyzed male datasets using matching parameters as in the current study. There was a 31% overlap of DEGs, (genes displayed in several timepoints were accounted only once for percentage calculation), with a total of 1294 up- and 1091 downregulated overlapping DEGs, respectively (adj. P-Value<0.05, Figure 5a-b). No overlapping genes were found in 1- and 2-wks groups in females and males (Figure 5c). Only 13 matching upregulated genes were identified between female LT and male 3wks groups, while in groups that displayed cachectic phenotype (female-HT and 4-wks-male), there were 1269 up- and 1091 downregulated genes matching between female and male LLC-bearing mice (Figure 5c). Amongst the top ten shared up- and downregulated genes for females and males, there were three (*Lcn2, Csf3r, Slfn4)* upregulated and one (*Tgtp2)* downregulated common genes. Downregulation of *Gbp* genes were exclusive to female LLC-bearing mice (Figure 5d-e).

To understand differences and similarities of CC-induced transcriptomic shifts in females and males, we compared dysregulated pathways (Figure 6). We found only one common downregulated pathway in 1wk groups and one common upregulated pathway in female LT and 3wks male groups. Most overlapping pathways were in the cachectic groups (female HT and male 4wks), with a total of 192 and 52 up-and downregulated respectively (Figure 6c). Considering the cachectic groups in both males (4wks) and females (HT) displayed the larger transcriptomic alterations, we focused subsequent analyses on 4wks and HT groups only. We next analyzed the top 20 disrupted pathways of each KEGG, Reactome, and WikiPathways, observing 37.5% of pathways were shared by cachectic females and males (Figure 6d, one repeated pathway was excluded). Amongst shared pathways, we identified upregulation of pathways associated with inflammation, metabolism of RNA, mRNA processing, autophagy, and exercise-induced circadian regulation, accompanied by downregulation of pathways associated with metabolism and mitochondrial systems (Figure 6d). We noted a more potent disruption of shared pathways in male compared to female mice, with male 4wks showing a more robust significance of the dysregulated pathways (Figure 6d, Adj. p-value<0.05). Unique dysregulated pathways for females and males are in Figure 6e-f and show female-unique upregulation of oxidative stress response, auto-degradation of the E3 ubiquitin ligase COP1, and proteasome among others. Upregulation of insulin signaling, cellular responses to stress, and metabolism of proteins among others, were unique to males (Supplementary Figure SF1). Additionally, distinct upregulation of *Interferon Pathways* including 15 genes (*Ube2l6, Isg15, Flnb, Ptpn6, Eif4g1, Eif4a3, Plcg1, Pias1, Abce1, Ptpn1, Kpna1, Ppm1b, Arih1, Kpnb1, and Ptpn11*) was noted in 4wks male (Data not shown, p=0.0001). Unique downregulated pathways to females displayed a relationship with mitochondrial pathways (Supplementary Figure SF1a), while male-only downregulated pathways included DNA replication and repair, estrogen signaling, and ribosomal pathways, in which 40% of genes in this pathway were dysregulated, including many Mitochondrial Ribosomal Proteins (MRBLs) and Ribosomal Proteins (RPSs) genes (Supplementary Figure SF1b).

**Figure 6.**
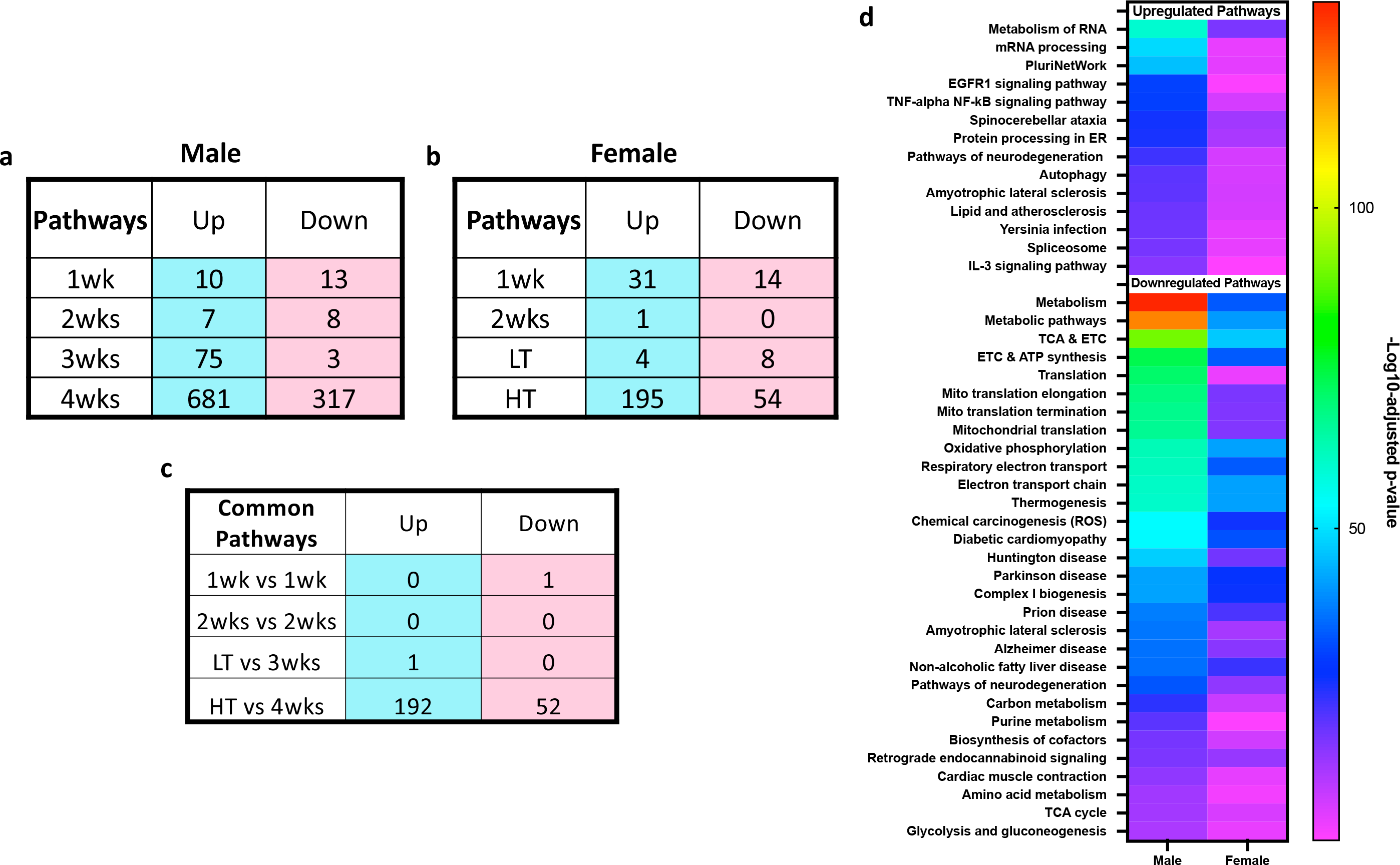
Female and Male Pathway Comparison. Number of dysregulated pathways in males (a), Number of dysregulated pathways in males (b), Matching Pathways in males and females (b) Top shared pathways in males and females (d).

After the observation of many mitochondrial-associated pathways being a dominant factor in the comparison of male and female mice undergoing CC, we inquired whether there were differences in MitoCarta matched genes in CC between biological sexes (Figure 7). Comparably with global gene expression, the most commonalities were found at the in the HT vs 4wks comparison, with 20 up- and 342 downregulated shared mitochondrial-associated genes (Figure 7a). Four genes displayed inverse expression profiling between biological sexes (Figure 7b). Figure 7c shows ~19% share of total upregulated mitochondrial-genes of females and males, highlighting the unique upregulation of apoptotic genes and mitochondrial gene expression-related genes in females, while fatty acid biosynthesis-related genes were uniquely upregulated in males. Downregulated mitochondrial genes displayed 54% shared genes between sexes (Figure 7d). Figure 7e shows distinction in Log_2_FC between the mitochondrial genes associated with the three top common pathways between biological sexes, highlighting a more prominent impact in males (higher Log_2_FC) compared to females.

**Figure 7.**
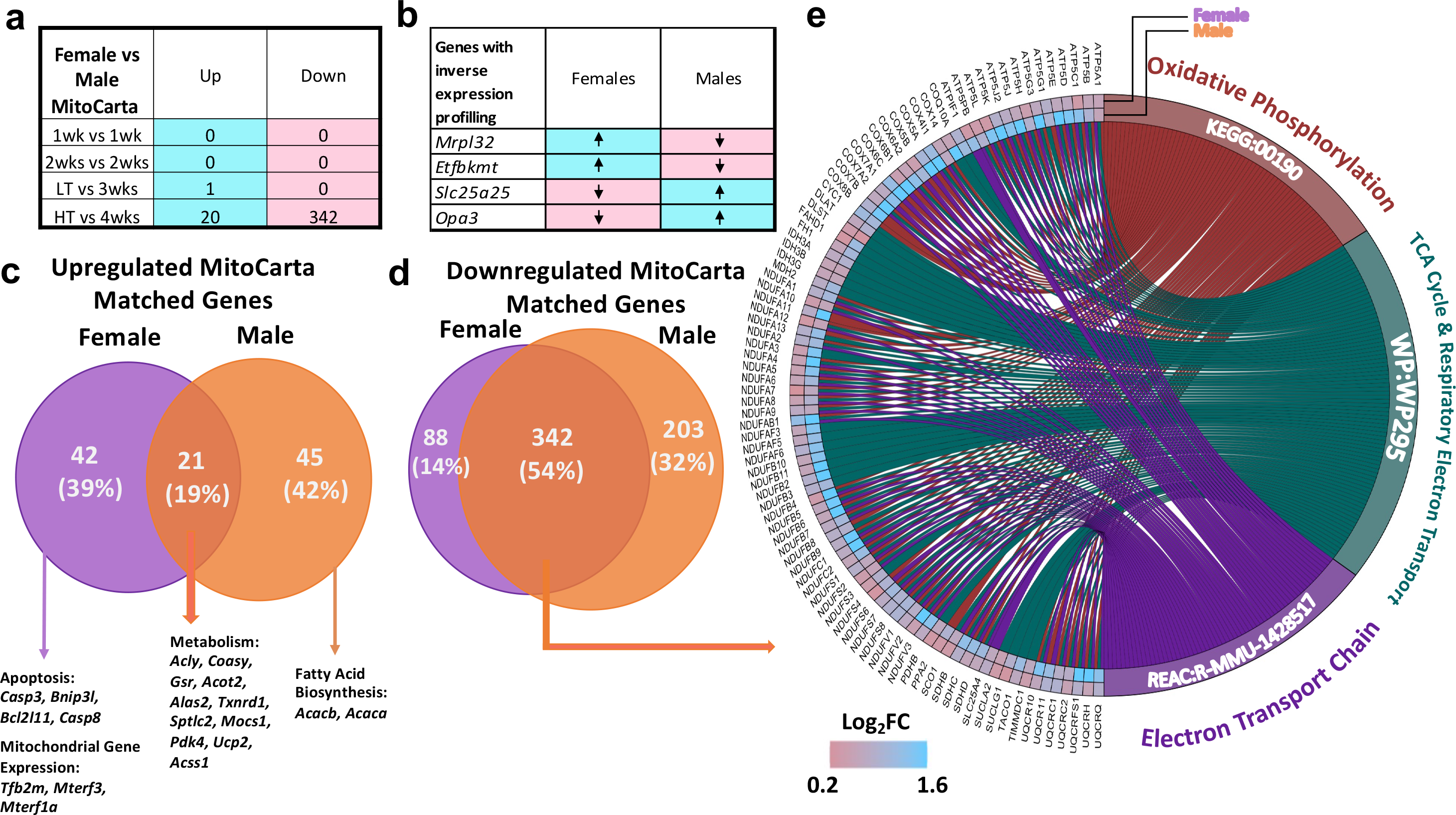
Biological dimorphism with focus on mitochondrial genes. Total number of shared mitochondrial genes in females and males at all timepoints (a). Inversed expression of four mitochondrial genes between biological sexes across all timepoints (b). Venn diagram of up- and down-regulated mitochondrial genes in females HT and males 4wks (c & d). Chord diagram display differences in Log_2_FC in common downregulated mitochondrial genes (e).

## Discussion

Effective treatments for CC remain lacking. Unfortunately, most preclinical studies have been historically performed in males, overlooking likely biological sex dimorphisms. Our group and others have shown differences in CC development between biological sexes (10, 17–21), demonstrating the necessity to study underlying mechanisms of CC unique to each biological sex. Here, we first evaluated the time-course of transcriptomic alterations in response to LLC-induced CC in female mice. We then directly compared cachexia-induced transcriptomic alterations between the current dataset in female LLC-bearing mice and reanalyzed data in male LLC-bearing mice (12) to compare between sex. In females, we observed muscle transcriptomic shifts one week following tumor allograft. Most importantly, major transcriptomic alterations occurred with onset of the cachectic phenotype despite relative preservation of gastrocnemius muscle mass in females. Further, our data shows disruption of important pathways involved in detriments to muscle function and health with cachectic development, including *cell structure, autoimmune system, protein ubiquitin, JAK-STAT pathways, and oxidative metabolism* in female mice. Cross-referencing female and male data (12), demonstrates only a ~33% overlap in DEGs of cachectic mice between biological sexes. Our data demonstrate unique downregulation of type-II-interferon genes amongst the top DEGs and pathways to female mice. Overall, data suggest a specific biological sex signature to CC progression.

### Overview of transcriptomic shifts in female mice undergoing LLC-induced cachexia

Our experimental model was successful in demonstrating key components of cachectic phenotype, including loss of fat mass, TA and soleus muscle wasting and splenomegaly. Transcriptional profiling of gastrocnemius muscle during different timepoints of tumor progression showed strong transcriptomic alteration one-week after cancer inoculation. Moreover, 83% of upregulated and 90% of downregulated DEGs were observed specifically in phenotypic cachexia. These findings are consistent with prior reports, confirming large transcriptomic shifts appear with onset of the cachectic phenotype (10, 12, 27). Corroborating the current study, prior research showed alterations at the transcriptomic levels in early and late stages of C26 colorectal and pancreatic ductal adenocarcinoma cancer-induced cachexia, despite mitigated muscle mass loss at early cachexia stage in females (10, 27). Altogether, the large degree of transcriptomic shifts occurs with the onset of cachexia.

### Consistent dysregulation across development of CC in females

To examine the DEGs further, we cross-referenced all DEGs between tumor-bearing groups. By cross-referencing we note 1wk and HT groups showed the most similarities, with 161 up- and 83 downregulated genes shared between groups, followed by LT and HT sharing 55 up- and 51 downregulated genes. Interestingly, only ten genes are consistently dysregulated across all timepoints, all of which were associated with downregulation of *Type II interferon signaling*; including upregulation of three type II interferon repressors with downregulation of seven genes associated with type II interferon activation. Furthermore, in reviewing the 10 most differentially expressed genes at each timepoint, several *Gbp* genes were downregulated across all timepoints. *Gbps-2,-3,-4,-5,-6* were amongst the ten most downregulated genes, many of which were downregulated in muscle across development of LLC-induced cachexia. *Gbps* are a GTPase family strongly induced by interferons and can contribute to cell survival through inhibiting apoptosis (28). One prior study shows downregulation of Type-II-interferon signaling in muscle of aged mice undergoing impaired regeneration (29), while another showed inefficient muscle regeneration in interferon null mice (30). Another recent study utilized single-cell RNA-seq to reveal upregulation in interferon-induced guanylate-binding protein through endothelial cells within skeletal muscle as a key player to aging-associated muscle loss (sarcopenia) (41) further suggesting the potential importance of interferon signaling in muscle atrophy. The role of Type II interferon signaling in muscle mass regulation is not yet fully understood, but this dataset supports a potential role for Type II interferon, specifically Gbps, in CC in females.

### Pathway analysis: biphasic transcriptomic alterations in females

We next examined functional ontology of CC-induced transcriptomic changes in female mice. Corroborating prior work in pancreatic cancer patients (31), pathways of *extracellular matrix (ECM) remodeling and cell structure* were upregulated in the 1wk group, including upregulation of genes associated with *collagen biosynthesis, deposition, focal adhesion* including multiple *Col-family genes*. Dysregulation in cell structure and induction of ECM remodeling, such as collagen deposition and fibrosis, have been associated with worsened prognosis for cachectic patients (31, 32), are observed in other tissues such as cardiac muscle (33), and are a hallmark of atrophy (15). Increased collagen deposition at the skeletal muscle endomysium is unique to cachectic patients, while cancer patients not presenting with cachexia do not exhibit collagen deposition (34). At 2wks, transcriptomic dysregulation was minimal, except upregulation of the *JAK-STAT* pathway, which persisted in the LT group, along with the upregulation of *inflammation and proteasome* pathways in LT and HT groups. JAK-STAT signaling is a known mediator of cachectic wasting (27, 35) associated with the acute phase response commonly noted in cachexia (27, 35).

Imbalanced protein turnover is a hallmark of muscle wasting (36). Corroborating this, *proteasome* was the strongest upregulated pathway in the HT group. The timing of this induction of the proteasomal system agrees with our prior findings (17) and ties induction of major catabolic signaling cascades with onset of the cachectic phenotype. Similarly, downregulation of multiple *immune system* associated genes and pathways was observed as early as one week following tumor allograft persisting throughout tumor development. Induction of the proteasome and dysregulation of immune-related functions found in the current study aligns with our prior report in males (12). However, *Type II Interferon signaling* downregulation (discussed above) was noted in female tumor-bearing mice only, displaying a biological sex dimorphism in CC. Interestingly, inflammation and immune system factors such as Nuclear Factor kB (NFkB) and Type II interferon are inversely regulated by ubiquitin proteasome systems, likely explaining concomitant downregulation of *immune-system* with increased *proteasome* activity. Yet, downregulation of Type II interferon preceded transcriptomic disruption in proteasome-associated genes in this dataset, raising the need for a deeper understanding of the role of type II interferon in muscle atrophy.

### Comparison of cachectic female DEGs and MitoCarta 3.0

We previously documented impaired mitochondrial health in multiple atrophy models, including LLC-induced cachexia (17, 19, 22, 37). Specifically, mitochondrial degeneration preceded atrophy in male mice undergoing either LLC-induced cachexia or disuse atrophy, while females largely protect mitochondrial health until onset of muscle atrophy (17, 19). Herein, we noted strong dysregulation in mitochondrial pathways including *oxidative phosphorylation, metabolic pathways, electron transport chain*, among others with the onset of cachexia itself. In fact, among the top 15 dysregulated pathways in HT mice, 11 were associated with energy metabolism. To further assess the extent of transcriptional disruption of mitochondrial genes we cross-referenced DEGs with the MitoCarta 3.0 showing an 18% match among all DEGs with 476 DEGs in HT mice from the MitoCarta (1158 genes in MitoCarta, 41% of MitoCarta genes). This data suggests large disruptions in mitochondrial gene expression correlates with onset of mitochondrial dysfunction (17) and cachexia. While mechanisms behind biological sex differences in cachexia remain largely elusive, the ability of female mice to protect muscle mitochondria during early stages of development may provide one such mechanism.

### Biological sex comparison in CC-induced transcriptomic alterations

Finally, we aimed to assess transcriptomic level biological sex differences in cachexia more quantitatively. Therefore, we returned to raw data from our male study and re-analyzed to match analysis parameters across studies. Most strikingly when comparing DEGs between cachectic male and female mice only ~1/3 of total DEGs were shared between sexes, meaning 2/3 were biological sex specific. This observation aligns with recent work in the KPC model of cachexia where in “late-stage” (i.e., cachectic) where a similar portion of DEGs were shared between biological sexes (10). Notably, within the 10 most up and downregulated genes for each biological sex only upregulation of *Lcn2, Csf3*r and *Slfn4* with downregulation of *Tgtp2* were observed across both sexes.

Similarly, across functional ontology only 37% of male pathways matched with females, while the same number represents ~98% of female pathways, demonstrating a larger transcriptomic disturbance in males. Among shared pathways, many were largely expected pathways associated with aspects of protein turnover and energy metabolism. Considering unique pathways, downregulation of multiple pathways associated with *DNA damage repair* and *ribosomal pathways* (including both mitochondrial ribosomal protein and ribosomal protein encoding genes) appeared specific to males, suggesting DNA maintenance and ribosomal activity play an important role in the protection in females but not male mice undergoing cachexia. This data is suggestive of the role of ribosomal activity in muscle mass regulation in cachexia discussed in previous studies showing reduced ribosomal capacity is associated loss of muscle mass (38) and is in accordance with mitochondrial detriments in males (19). Females did not display dysregulation of ribosomal pathways, accompanied by preservation in gastrocnemius mass, while the opposite was noted in males, suggestive of a protective role of muscle mass in maintenance of ribosomal function.

Additionally, while female mice show consistent downregulation of interferon type II associated genes in multiple timepoints of CC development, males show upregulation of interferon-genes at 3- and 4-weeks following tumor allograft, raising possibility of a novel interferon role in cancer-induced atrophy. Meanwhile, females displayed unique upregulation of *exercise-induced circadian regulation*, including *Pura*, and *Ncoa4*, which are associated with regulation of DNA replication, preventing inappropriate DNA synthesis and replication stress (39). Moreso, females showed upregulation of *Oxidative Stress Response*, including important antioxidant-associated genes including *Sod1* and *Cat* (40). Upregulation in genes associated with aid to DNA replication and antioxidant response may partially explain relative protection to muscle and mitochondrial health noted in females (17) compared to males undergoing CC. While dysfunctional mitochondria (19) precede prominent alterations in mitochondrial pathways in males (12), concomitant transcriptomic and functional mitochondrial alterations (17) are delayed in female skeletal muscle. Altogether, data suggests key biological sex dimorphisms in functional and intrinsic skeletal muscle alteration and therefore sex-specific mechanisms of cachexia.

### Perspectives and Significance

Despite the potentially critical importance of biological dimorphisms withing health and disease, the majority of pre-clinical work in CC is still performed in male specimens. This study focused on exploring the female transcriptomic alteration in response to tumor presence and dissect distinctions when compared to male mice. This study contributes to a larger body of evidence to support the existence of biological sex differences in the development of cancer-induced cachexia. Within our study approximately two-thirds of all DE genes were biological sex specific. Furthermore, we unveil the downregulation of interferon-associated genes to be unique to female, concomitant to protection of skeletal muscle mass in the presence of systemic cachectic phenotype. Males on the other hand show elevated interferon signaling, along with marked impairments in energy metabolism pathways and loss of muscle mass. This finding represents a new potential therapeutic target for mitigation of poor CC outcomes through strategies to reduce interferon signaling in skeletal muscle environment. Overall, our data strongly suggest sex specific mechanisms of CC which must be considered in further understanding this debilitating condition and developing appropriate and efficacious therapeutic modalities.

### Conclusion

Overall, our study adds to the growing evidence demonstrating biological sex differences in development of CC. Here we showed two large disruptions in transcriptome in female LLC tumor-bearing mice; one associated with alteration to the extracellular matrix, and a second accompanied by development of the cachectic phenotype affecting overall skeletal muscle protein and energy metabolism in parallel with functional decrements in mitochondria and muscle health. Furthermore, alterations to Type II Interferon signaling appears to be a signature of CC development in females, accompanied by preserved gastrocnemius mass not noted in males (Figure 8). By comparing data in female mice with our prior work in male LLC-bearing mice, we reveal only ~one-third of DEGs are shared between biological sex, strongly indicating biological sex dimorphism in transcriptomic response to cachexia. This study is limited to preclinical analysis and use of a single cancer type. Herein, we have demonstrated novel transcriptomic alterations to skeletal muscle in female mice undergoing LLC-induced cachexia distinct from those observed in males. These data strongly suggest biological sex specific mechanisms in CC which need to be considered in developing effective therapeutic approaches to prevent and reverse this condition.

**Figure 8.**
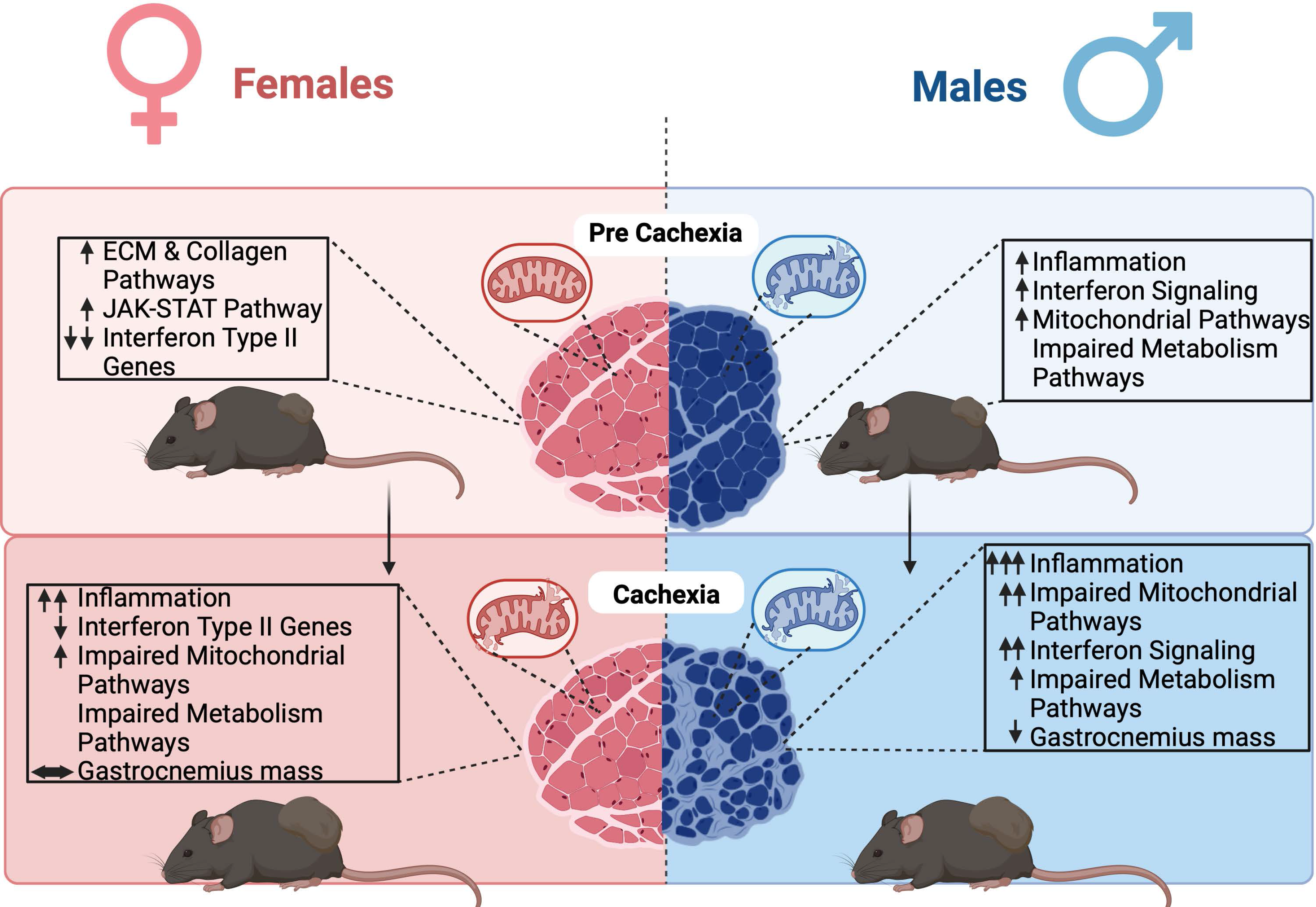
Summary figure. Females and males display different timeline transcriptomic responses to tumor development and cancer cachexia. Females display an early dysregulation in extracellular matrix remodeling and cellular structure associated pathways followed by increase in JAK-STAT pathways associated gene expression. Interesting, besides maintenance of gastrocnemius muscle mass, downregulation of interferon type II genes is unique to females through cancer cachexia development. Delayed transcriptomic changes in mitochondrial pathways match functional mitochondrial impairments previously reported. Males show upregulation inflammation pathways, and interferon signaling genes at pre-cachectic and cachectic states. Males show early disruption in mitochondrial function that precedes loss of gastrocnemius mass and major transcriptomic changes in mitochondrial genes. Invert expression profiling of interferon-associated genes between biological sexes represent a transcriptomic signature to each biological sex in response to cancer cachexia development and suggest a potential target for muscle loss during cancer-induced cachexia.

## Supporting information

Supplementary Information Description

## Declarations

### Ethics approval and consent to participate

All animal methods were approved by the Institutional Animal Care and Use Committee of the University of Arkansas.

### Consent for publication

Not applicable.

### Availability of data and materials

Raw sequencing data will be deposited in a publicly available database upon publication.

### Author’s Contributions

Manuscript was written by FMS with support from all authors (ARC, ERS, TAW, NPG, KAM, RJIII, SL, MRC). NPG designed experimental and analytical approach. Animal experiments were conducted by MRC, SL, TAW and NPG. RNA sequencing analysis and presentation were done by FMS with support from KAM and NPG. RJIII contributed to chord diagram presented in this paper. All authors approved the final manuscript.

## Acknowledgments

Thank you to Mr. Kevin Greene and Dr. Aaron Caldwell for their aid in coding and cross-referencing.

## Author’s Information

^1^Cachexia Research Laboratory, Exercise Science Research Center, Department of Health, Human Performance and Recreation, University of Arkansas, Fayetteville, AR

^2^Exercise Muscle Biology Laboratory, Exercise Science Research Center, Department of Health, Human Performance and Recreation, University of Arkansas, Fayetteville, AR

^3^Molecular Muscle Mass Regulation Laboratory, Exercise Science Research Center, Department of Health, Human Performance and Recreation, University of Arkansas, Fayetteville, AR

