## Supplementary figures and images for "The Time-Course of Cancer Cachexia Onset Reveals Biphasic Transcriptional Disruptions in Female Skeletal Muscle Distinct from Males"

### Supplementary Figure SF1

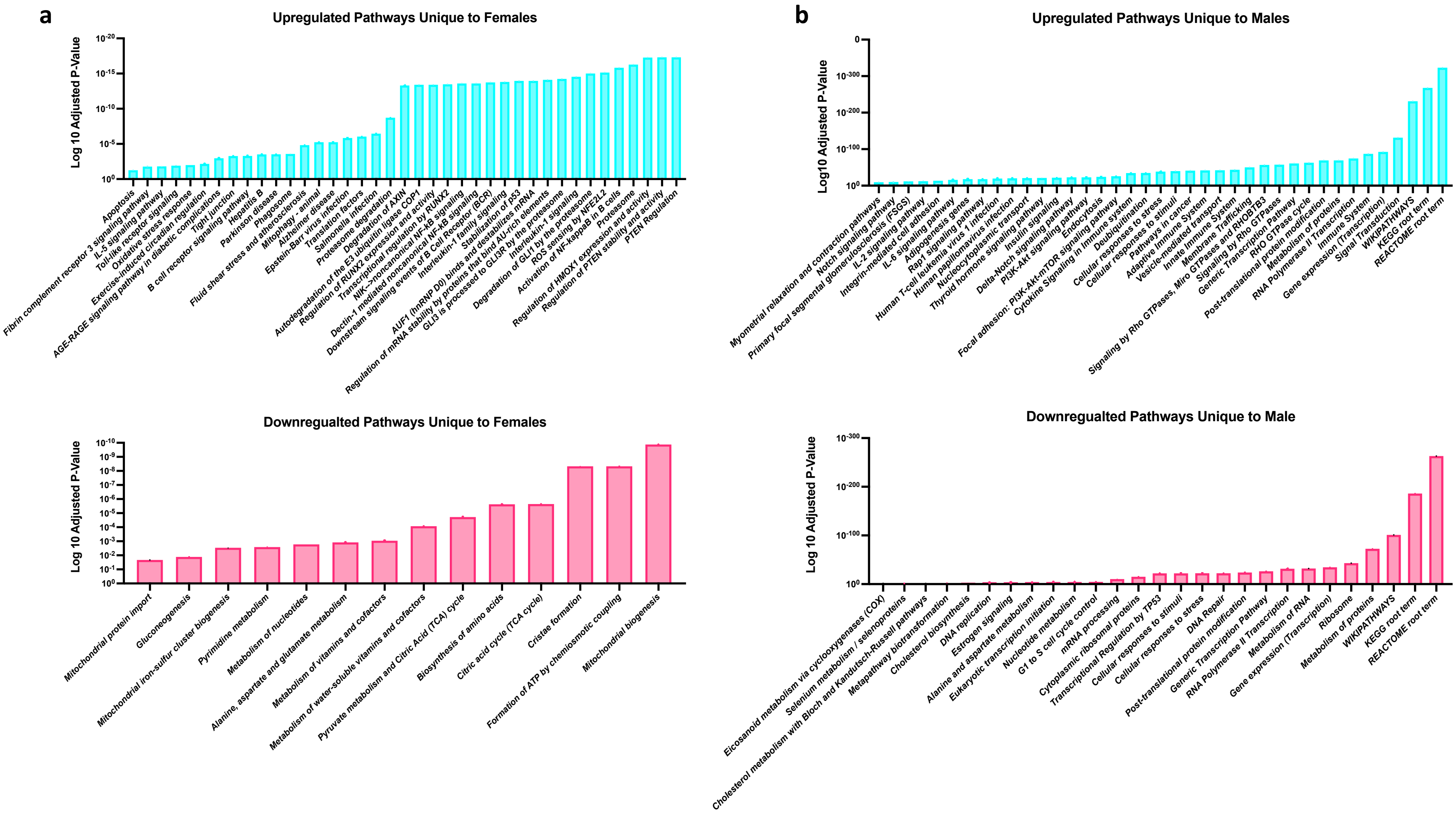
