## Supplementary Information Description for "The Time-Course of Cancer Cachexia Onset Reveals Biphasic Transcriptional Disruptions in Female Skeletal Muscle Distinct from Males"

Supporting Information & Supplementary Figure

**Supplementary Figure SF1.** Top unique to female dysregulated pathways (a), Top unique to male dysregulated pathways (b). Top 20 of each Kegg, Reactome, and WikiPathways. Adjusted P-value<0.05.

**Supporting Information S1.** Female DESeq data output.

**Supporting Information S2.** Female Pathway Analysis.

**Supporting Information S3.** Female DEGs vs Mitocarta.

**Supporting Information S4.** Male DEseq data output.

**Supporting Information S5.** Female vs Male DESeq.

**Supporting Information S6.** Male Pathway Analysis.

**Supporting Information S7.** Female vs Male Pathway Comparison.

**Supporting Information S8.** Male DEGs vs Mitocarta.

**Supporting Information S9.** Female vs Male Mitocarta Comparison.
